# Trait Evolution on Phylogenetic Networks

**DOI:** 10.1101/023986

**Authors:** Dwueng-Chwuan Jhwueng, Brian C. O’Meara

**Affiliations:** Department of Statistics, Feng-Chia University No. 100, Wenhwa Rd., Seatwen, Taichung, Taiwan 40724, R.O.C.; Department of Ecology and Evolutionary Biology, University of Tennessee, Knoxville, TN, 37996-1610, USA

**Keywords:** hybridization, gene flow, phylogenetic comparative methods, Brownian motion, network

## Abstract

Species may evolve on a reticulate network due to hybridization or other gene flow rather than on a strictly bifurcating tree, but comparative methods to deal with trait evolution on a network are lacking. We create such a method, which uses a Brownian motion model. Our method seeks to separately or jointly detect a bias in trait value coming from hybridization (*β*) and a burst of variation at the time of hybridization (*v*_*H*_) associated with the hybridization event, as well as traditional Brownian motion parameters of ancestral state (*μ*) and rate of evolution (*σ*^2^) of Brownian motion, as well as measurement error of the tips (SE). We test the method with extensive simulations. We also apply the model to two empirical examples, cichlid body size and *Nicotiana* drought tolerance, and find substantial measurement error and a hint that hybrids have greater drought tolerance in the latter case. The new methods are available in CRAN R package *BMhyd*.

---

Various comparative methods have been proposed to deal with the fact of non-independence of species due to shared history on a phylogenetic tree (Felsenstein 1985, 2008; Cheverud et al.1985; Grafen 1989; Gittleman and Kot 1990; Hansen 1997; Lynch 1991; Housworth et al. 2004; Butler and King 2004; O’Meara et al. 2006; Hansen et al. 2008; Beaulieu et al. 2012) as well as many other problems, ranging from ancestral state estimation (Schluter et al. 1997) to estimating the effect of traits on diversification rates (Maddison et al. 2007) to predicting extinction risk (Cardillo et al. 2006). However, hybridization, a fairly common process (Mallet 2005, 2007; Riesberg 2006), results in species being related in a phylogenetic network rather than a tree (Arnold 1996; Doolittle 1999; Otto and Whitton 2000; Linder and Rieseberg 2004; Huson et al. 2010; Nakhleh 2011), and this reality is not yet accomodated by existing comparative methods. Due to the importance of hybridization as a process, numerous methods have been developed to infer phylogenetic reticulate networks (for simplicity, we refer to these as “networks”) rather than trees (e.g., Sang and Zhong 2000; Weigel et al. 2002; Bryand and Moulton 2004; Moret et al. 2004; Huson and Bryant 2006; Joly et al. 2009; Kubatko 2009; Meng and Kubatko 2009; Wang et al., 2013; Willson, 2013; Wu, 2013). We thus stand at a point where phylogenetic networks will increasingly be inferred but we lack comparative methods to properly use this information.

A species formed as a hybrid of two parental species can differ from its parents in important ways. Due to transgressive segregation (Rieseberg et al. 1999), hybrids may have trait values outside those of their parental species. If we consider species’ trait values evolving through time in a Brownian motion random walk, this transgressive segregation can be modeled in a few different ways. For example, if hybrids are on average 10% larger than their parent species, this could be modeled by a shift in mean trait value associated with hybridization. If processes like transgressive segregation lead to difference from parents but with no consistent trend in direction across many hybrid events, this could be modeled as a burst of variation at the time of the hybrid event. Hybrids may also evolve at different rates than parental species, especially if they are formed from polyploidization (Ainouche et al. 2008). This could be reflected in a different rate parameter for hybrid species than for non-hybrids. Finally, hybrids may not be formed equally from both parental species. For example, one “hybrid” species may have formed through regular allopatric speciation of a single species, plus a few genes introgressed from a neighboring species. If a phenotypic trait value represents the additive result of multiple quantitative loci, an appropriate model would treat the hybrid trait mean as being a weighted average of the two parental species’ means, with the weighting based on the relative genetic contributions of each parent.

In this work, we propose a new comparative method to study trait evolution under phylogenetic networks. This method allows for estimation of traditional evolutionary parameters such as rate and overall mean under a Brownian motion model while also allowing investigation of trait evolution occurring as a result of the hybridization process. We test our model with simulations and investigate two empirical datasets of cichlid and *Nicotiana* (tobacco and relatives).

## METHODS

### Brownian Motion for Trait Evolution

Brownian motion (BM) is a general model for unbounded continuous trait evolution commonly used in phylogenetics (Felsenstein 1985). Biologists often incorrectly believe this is only a model for traits evolving under genetic drift, but in fact a variety of biological mechanisms can lead to this same model, such as selection towards an optimum that changes due to multiple factors through time, drift-mutation balance, an evolutionary trend, as well as pure genetic drift (Hansen & Martins 1996). Under Brownian motion, the variance of a trait is proportional to evolutionary time multiplied by the rate of evolution, *σ*^2^. Therefore, given a phylogenetic tree the covariance among species can be represented by the shared branched length on the phylogenetic tree. Figure 1 shows a three-taxon phylogenetic network with gene flow.

**Figure 1:**
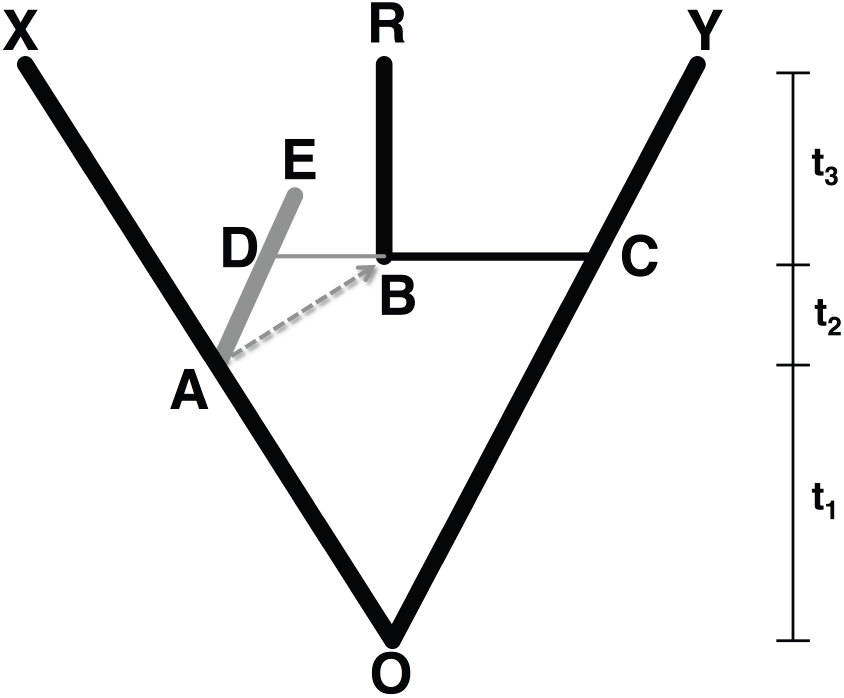
Three taxon network with extant species X, R and Y. Species C and species D are the immediate parents of species B; in this example, it gets 75% of its genes from C, 25% from D. This network can be thought of as a tree (the black edges) combined with gene flow from A via D into B (the gray edges). D’s descendant, E, is not in the final tree that is used (X,(R,Y)) either through extinction or not being sampled. Thus, even though gene flow must occur between coeval species, the effective path of gene flow from the X lineage to the R lineage occurs starting at time *t*_1_, not time *t*_1_ + *t*_2_, the actual origin of the hybrid: the changes occurring on the X branch in the *t*_2_ time interval cannot be shared with the hybrid B. The dashed arrow starting at time *t*_1_ and ended at time *t*_1_ + *t*_2_ thus indicates the effective gene flow from A to B.

This represents a scenario where at time *t*_1_, there was a speciation event: one branch led to X, and the other led to a species D that eventually went extinct (E) or was otherwise unsampled in this analysis. However, at time *t*_1_ + *t*_2_, species D exchanged genes with the species at C to form a hybrid species, B, which survived to be sampled species R. Though gene flow only occurs between taxa occurring at the same point in time (D and C), due to extinction it can look like flow forward in time: the shared history of R with X is not from when D exchanged genes (time *t*_1_ + *t*_2_) but earlier, time *t*_1_: changes on the A to D branch are not shared between X and R. Thus, the dashed line shows the effective path leading to the covariance of the observed tips, rather than the path from A to D to B and thus to R. The corresponding variance-covariance matrix for the extant species X, R, Y for the tree model (black edges only) is given by the matrix **V**_***s***_ as following

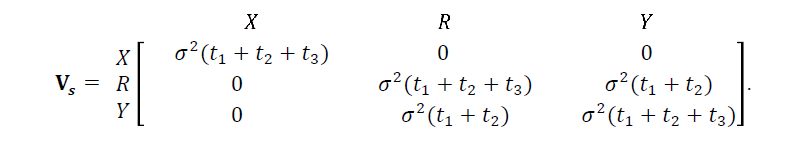

Now consider the trait evolution includes the gene flow (dashed arrow). Let the trait value of the root state *O* be *μ*. By assuming species evolve under Brownian motion (variables are measured in log scale in comparative analysis), the trait values at species D and species C are *μ*_*D*_ = *μ* + ε_*D*_ and *μ*_*C*_ = *μ* + ε_*C*_, respectively, where *ε*_*D*_ and *ε*_*C*_ can be regarded as error terms that follow a normal distribution with zero mean and variance *σ*^2^(*t*_1_ + *t*_2_). Under our model, the hybrid species *B*, at the moment of hybridization, assumes the value *μ*_*B*_, defined as

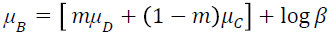

which follows a normal distribution with mean *μ* + log *β* and variance *σ*^2^(*t*_1_ + *t*_2_). The parameter *m* measures the proportion of the hybrid trait value inherited from parent D while 1 − *m* measures the proportion the hybrid inherits from parent C (*m* is bounded between zero and one). If the hybrid species is formed mostly from individuals of species D, with only some gene flow from species C, then for a polygenic quantitative trait, *m* might be much closer to one. In particular, *m* = 0.5 indicates inheriting the trait equally from both parents. In the absence of information to the contrary, 0.5 represents a reasonable value to use. The parameter *β* governs the possible bias in trait value as a result of hybridization. If there is a bias that leads to greater fitness, this is often called heterosis or hybrid vigor; if there is a bias that leads to lower fitness, this may be called outbreeding depression. Here we care about trait values, not their fitness effects, but hybrid means may be thought of in the same way, in that they may be something other than the average of their parents. For example, if there exist widespread heterosis, with hybrids being on average 20% larger than their parent species, *β* would be 1.2. The natural lower bound for *β* is zero and the upper bound is arbitrary; a value of 1 indicates that the hybrid is just a weighted average of its parents. Brownian motion assumes that an increase or decrease by a certain amount has the same probability regardless of a trait value. This is often not the case for raw measurements: an increase or decrease of mass by 1 kg over a million years is far likelier for an elephant species than for a mouse species. However, they both might be equally likely to increase or decrease their mass by 1%. It is thus typical to log transform raw values to meet this assumption. Therefore, we add the log parameter, log *β*, to represent log scale bias for the hybrid at formation. Here we assume that *μ*_*A*_, μ_*B*_ and *μ*_*C*_ were in log scale already (i.e. the representation in raw scale is 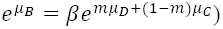. To model a process like transgressive segregation, where a hybrid can deviate from the range of parental values but without a particular bias, non-negative variance *v*_*H*_ is added to lengthen the hybrid branch, equivalent to adding a burst of variation due to the hybridization event. Therefore, we have *Var*(*X*) = *Var*(*Y*) = *σ*^2^(*t*_1_ + *t*_2_ + *t*_3_) and *Var*(*R*) = *σ*^2^(*t*_1_ + *t*_2_ + *t*_3_) + *v*_*H*_. To allow hybrid species to have different rates of evolution than non-hybrid species, one would just require modifying this variance to be 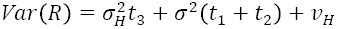, where the new term 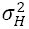 is the rate of evolution in hybrids. We currently limit the model to the one where the hybrid and non-hybrid species have the same rate(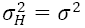). The corresponding variance-covariance matrix for the species *X*, *R*, *Y* for the network model under the assumption of the BM process for trait evolution is given by the matrix ***V***_***R***_ as following

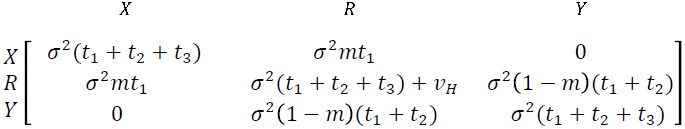

Furthermore, measurement error can be substantial, and it has the effect, if ignored, of leading to larger estimates of rates on tip branches. We deal with this (following a suggestion in O’Meara et al. (2006)) by adding a parameter, SE, to the diagonals of ***V***_***R***_ to represent measurement error. The final ***V***_***R***_ thus becomes:

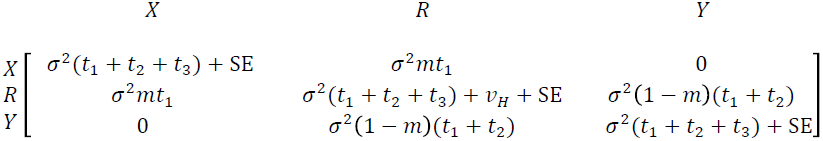

Given traits *y*_1_, *y*_2_, …, *y*_*n*_ (in log scale) for *n* species some of which are hybrids, the column vector ***Y*** = (*y*_1_, *y*_2_, …, *y*_*n*_)^*T*^ can be treated as a multivariate normal random variable given an assumption of Brownian motion. i.e. ***Y*** ∼ *MVN*(***μ***, ***V***_***R***_) where ***μ*** = (*μ*_1_, *μ*_2_, …, *μ*_*n*_)^*T*^ and *μ*_*i*_ = *μ* or *μ* + log *β* is the mean for usual species or the mean for hybrid species, respectively, and **V**_***R***_ is calculated as above given the tree (with branch lengths) ***T***, structure of gene flow ***D***_*t*_d_,*t*_*r*_,*m*_, and parameters μ, *σ*^2^, *v*_*H*_, and *β*. The negative log likelihood function given these is

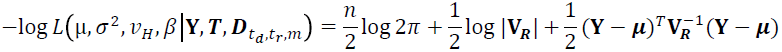

where |**V**_***R***_| is the determinant of **V**_***R***_ and (**Y** − ***μ***)^*T*^ is the transpose of (**Y** − ***μ***). **D**_*t*_*d*_,*t*_*r*_,*m*_ contains information on gene flow. It must be provided by the user, and indicates donors, recipients, and the time of hybridization events.

### Numerical problems

It is known that the variance covariance matrix of the phylogeny can be ill-conditioned: the matrix can be effectively singular, which makes dealing with its inverse, as required to calculate the likelihood, numerically difficult. Anecdotally, this seems to occur more frequently for phylogenetic networks than trees. One approach would be to just prohibit analyses if this is the case. This is practically problematic: a biologist spending years gathering data and a tree could find her or his analysis stymied just due to the structure of the network. Instead, we opted for an approximate solution a user may choose to apply (though the default is not to do this). We evaluate the condition by taking the variance covariance matrix and calculating the determinant of it, of it multiplied by 1000, and of it multiplied by 0.0001. If all three determinants are positive, the matrix is more likely to be well-conditioned, no matter the parameter estimates (which, in the case of *σ*^2^, *v*_*H*_ and SE, may change the magnitude of entries). If it fails this test, we lengthen terminal branch lengths slightly and try again, repeating until the maximum number of tries (by default, 100, causing the terminal branch lengths, likely to be in units of millions of years, to be increased by just 0.001). If it is still ill-conditioned, the analysis aborts; otherwise, it continues, but will only return an approximate answer. This adjustment is done to the original tree, before fitting parameters; regardless of whether that was done, the matrix may also be poorly conditioned for some parameter combinations, resulting in numerical errors in calculation of likelihood. During a run, the log of the condition number of the final variance covariance matrix is measured and compared to a set precision; if this value exceeds the precision, numerical problems may ensue. We approximate what the likelihood would be given the problematic variance covariance matrix by calculating the likelihood with a series of better conditioned ones (by decreasing the magnitude of off diagonal elements while increasing the magnitude of diagonal elements), and then predicting the likelihood with no such transformation from a smooth spline extrapolation of the likelihood from matrices with decreasing magnitude of transformations. This approximation is turned off by default, but can be helpful if the maximum likelihood estimates occur in a problematic region. The transformation of the mean vector proceeds similarly.

### Assessing the general performance of the model

We assessed the performance of our model, varying (i) the number of non-hybrid taxa (30 or 100), (ii) the number of hybrids (1, 5, or 10), (iii) the proportion of flow of hybrid inherited from the parents (all from one parent, 10% from one and 90% from the other, or equally from both parents), (iv) the structure of hybridization, (v) the value of *β* (0.1, 1, or 10), and the value of *v*_*H*_ (0, 10, or 100). For each replicate we simulated a bifurcating tree of 30 or 100 taxa using TreeSim (Stadler 2009), with a birth rate of 1, death rate of 0.5, sampling frequency of 0.5, and tree height of 50. We then added 1, 5, or 10 species of hybrid ancestry to the tree in one of two ways. The first was to attach those taxa randomly around the tree, but forcing each to arise from its own hybridization event: that is, no hybrid species subsequently speciated. The second was to have just one hybridization event on the tree, and have this lead to the observed number of species of hybrid origin through speciation of the original hybrid. Different simulations differed in the ratio of genes coming from the two parents: it could be all from one parent (a flow of 0: this is equivalent to a tree model); 10% from one parent and 90% from the other; or 50% from each parent. Other parameter values fixed in the model were *μ* = 1, σ = 0.01, and SE = 0. All simulations were carried out by the *BMhyd* package. After the runs were done, we looked at deviation (absolute value of the difference between the observed and true parameter value, divided by the true parameter value (if it is nonzero)). We regressed this for each parameter of interest against the values used in our simulation: tree type, flow magnitude, number of non-hybrid taxa, number of hybrid taxa, and how a summary of the parameters could be calculated (using only the best of four models, doing model-averaging across all four models, or doing model-averaging across the models with *v*_*H*_ fixed). The two 3-state values (flow and summary approach) were converted into two binary variables each. The package MuMIn (Barton 2015) was used to estimate the Akaike importance and coefficients for each of these parameters. The goal of this was to allow us to focus figures and discussion on the simulation parameters that had a strong effect on the relative error, rather than try to plot the seven dimensions of variation used in the simulations.

### Empirical test cases

Simulations are essential in examining a new method to verify that it is working well enough in choosing models and estimating parameters. However, it can also be useful to run empirical datasets, both to make new discoveries using a new method and to verify that a method operates smoothly on real, messy data. Unfortunately, as the true model or parameter estimates are not known from empirical data, information about accuracy may only come from the simulations, but empirical results can show other problems. In this paper, we look at hybridization in two examples: cichlid body size evolution, using a network from Kobmüller et al. (2007) and data from FishBase (Froese and Pauly 2010), and tobacco and relatives, using a network from Chase et al. (2003) and drought tolerance data from Komori et al. (2000).

### Cichlids

Cichlids are notorious for widespread hybridization; their phylogeny is difficult due to presumed hybrid origin or ongoing gene flow. In fact, some may be going extinct due to merging through hybridization (Rhymer and Simberloff 1996). They thus reflect a good test case for this method. Kobmüller et al (2007) developed a phylogeny for cichlids which, importantly, included information about hybrid species and their presumed direction of ancestry. To replicate their tree, we downloaded their sequences from GenBank (Benson et al. 2005). The sequences were aligned by MAFFT (Katoh and Standley 2013) and subsequently inspected in Mesquite (Maddison and Maddison 2011) to trim ends of sequences for only a small subset of taxa. A backbone constraint was made from Kobmüller et al (2007)’s overall hybrid tree in Mesquite and used in all subsequent searches. The aligned sequences were then analyzed under the software PAUP (Swofford 2003) for parsimony tree search. The searched best-rooted tree was taken and was used to set to likelihood of GTR + Γ substitution model with clock. We filtered for best under this model and then did a search limited to 10 hours to resolve branch lengths. Trait values (total body length of cichlids) were collected from FishBase (Froese and Pauly 2010) using the R package rfishbase (Boettiger et al. 2012).

In Kobmüller et al. (2007), the cichlid data contains 27 species where five species are putative hybrids. Three of the five hybrid species (*Neolamprologus wauthioni, Lamprologus speciosus*, and *Neolamprologus fasciatus*) are inferred to have arisen due to mating between extant species and two of them (*Lamprologus meleagris, Neolamprologus multifasciatus*) are inferred to be formed as a result of hybridization between extinct lineages. As in Fig. 1, this results in the gene flow appearing to be forward in time from at least one relative of the parental species.

### Nicotiana

This group contains tobacco and relatives. Their relationships were long suspected to be reticulate (Godspeed 1954), and this was supported by Chase et al. (2003) in a work based on internal transcribed spacer region (ITS) and in situ hybridization. We followed the same procedure as for the cichlid dataset in returning a chronogram, except that we did not use Chase et al.’s parsimony trees as constraints. The crown age was set to 15.3 MY, following Clarkson et al. (2005). Taxa of hybrid origin and the placement of hybridization events were pulled from Chase et al.’s results; timing of events came from branch lengths on the chronogram, where the donor and recipient times were set to be equal (thus, no postulate of extinct intermediate hybrid parents) and to occur at the origin of the hybrid taxon. The relative seedling growth under mannitol treatment dataset from Komori et al. (2000) was extracted from their Table 2. We note that this number is a proportion, thus not quite meeting the expectations of Brownian motion (unbounded traits); we log transformed it, but this is still an imperfect fix. We used iPlant TNRS (Boyle et al. 2013) to convert the taxon names from both datasets to the same taxonomy, and Geiger (Harmon et al. 2008) to prune the tree and data to the same taxon set. Figure 2 represent the evolutionary tree and the gene flow for both cichlid and tobacco. The donor-recipient relationship among the hybrids and their parents in the cichlid and *Nicotiana* datasets can be found in Supplemental material.

**Figure 2:**
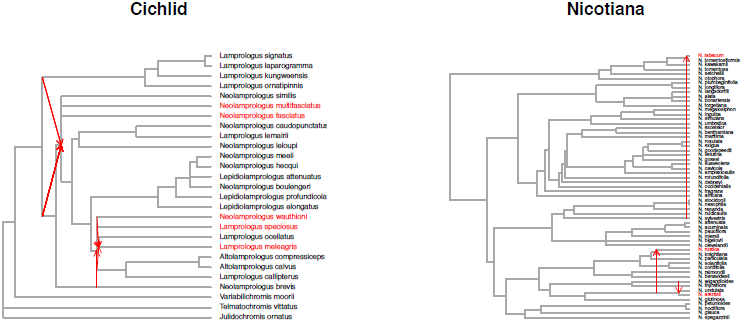
The empirical networks. Species in red are of putative hybrid origin. Red arrows show movement from a parent to a new hybrid lineage; in cases where only one arrow is shown leading to a lineage, it is because the hybrid lineage comes from its sister species on the tree plus the source of the arrow. Arrows appearing to move forward in time show transfer via an unsampled lineage (see explanation on Fig. 1).

### Model Selection and Parameter Estimation

We tried four different models for each empirical dataset. All fix the gene flow *m* = 0.5 and allow *μ*, *σ*^2^, and SE to be optimized. They differ in the settings for mean change in the hybrid *β* and the hybrid variation at formation *v*_*H*_. Model 1 fixes *β* at 1 but allows *v*_*H*_ to vary; Model 2 allows *β* to vary but fixes *v*_*H*_ at 0; model 3 fixes *β* at 1 and *v*_*H*_ at 0 and model 4 allows both to vary. We fit those models to both cichlid and tobacco datasets.

### Adaptive confidence intervals sampling

Uncertainty in parameter estimates can be substantial. One way of estimating this can be looking at the curvature of the surface at the maximum likelihood optimum, but this is known to be problematic when the likelihood function is not regular (Pawitan 2013). A different approach, advanced by (Edwards 1992) is to look at a confidence region of all points that generate a log likelihood within a certain range (often, set to be a delta of 2 log likelihood units) of the maximum likelihood. One approach to calculate this would be to vary each parameter on its own while holding the others constant. This is convenient and fast to implement, but can result in artificially small confidence intervals. For example, if two parameters *a* and *b* covary such that the likelihood is the same as long as *α* = 0.7*b*, the likelihood changing just *a* or just *b* would drop off very quickly, but there’s a ridge containing a wide array of *a* and *b* values that would not affect the likelihood. Thus, we chose to examine varying all parameters at once, so that if there is a ridge or other structure for the likelihood surface we do not overestimate our certainty. While there are many algorithms to find the peak of a surface, there are fewer to find the entirety of a region two log likelihood units below the peak. We thus developed a Monte Carlo method to estimate this. We start by simulating points using a multivariate uniform centered on the maximum likelihood estimates. The likelihood at each of these points is calculated. The algorithm periodically checks to make sure half the points are within the region and half are outside. If too many are within the cutoff of the peak likelihood, there is not good enough sampling of the boundaries of the confidence region and the sampling width is increased; if there are too many that have values too far from the optimal likelihood, the sampling width decreases. For a given parameter value, we thus calculate the likelihood over a range of values for the other parameters, giving a more realistic, less conservative confidence interval. Note, however, that this merely examines uncertainty due to flatness of the likelihood surface: there can be substantial additional sources of uncertainty from tree topology or branch length uncertainty, problems with measurements beyond what a fixed measurement error can address, or other issues.

### Software and Data

We have implemented this model in R (R team 2015) in the *BMhyd* package (on CRAN). It is open source, and includes functions for fitting models on networks, visualizing gene flow on networks, and even simulating random networks. It uses functions or code from *Geiger* (Harmon et al. 2008), *phytools* (Revell 2012), TreeSim (Stadler 2014), ape (Paradis 2004). All relevant R code and data files in this work can be found at Dryad Digital Repository

## RESULTS

### Simulation for Assessing General Performance of Models

Overall we were not pleased with performance: looking at just the simulations with the best chance for good results (flow rate of 0.5, 100 non-hybrid taxa, 10 hybrid taxa) there was often quite poor correlation between the estimates of parameters *v*_*H*_ and *β* and the true values (SE, *μ*, and *σ*^2^ performed far better, but they are not interesting parts of our model). Experimenting with the results suggested that model averaging or just taking the parameter estimates from the best model had about the same performance; using models where only *β* varied performed better than models where *v*_*H*_ varied. We investigated this, and other elements of the simulation, by doing a regression where we found importance and coefficients of parameters by dredging (using all subsets of the global model, and using Akaike weights to calculate importance). This suggested that choosing model-averaged or best model only parameter estimates did not matter much (importance of 0.27) but using models with *v*_*H*_ fixed was important for estimating *v*_*H*_ (importance of 1.00), less so for *β* (importance of 0.27) [and important for the other three free parameters], but the sign of the coefficient, negative for both *v*_*H*_ and *β*, suggested that limiting models to those with *v*_*H*_ fixed reduced error in both *β* and, oddly, *v*_*H*_. Tree type was not important. This surprised us: for ten hybrid taxa, having them the outcome of ten independent hybrid events should give much more information about the hybridization process than ten hybrids descended from one event in the past. However, inspecting the parameter estimates coming from trees with each simulation scenario also suggested they performed very similarly. Summaries later in the paper thus merge trees of both types. Flow and number of hybrid and nonhybrid taxa were found to be important.

Model-averaged parameter estimates for the most relevant parameters *β* and *v*_*H*_ in simulations are shown in Fig.3; following the information on importance, above, we grouped results regardless of hybridization type, and only show results with equal flow from both parent species (though results were similar with flow of 10% from one parent and 90% from the other, as well as, oddly, flow from only one parent). The estimates for all parameters, using model averages from the set of all models and from the set of models that had *v*_*H*_ fixed at zero, are shown in Supp Fig 1.

**Figure 3 caption:**
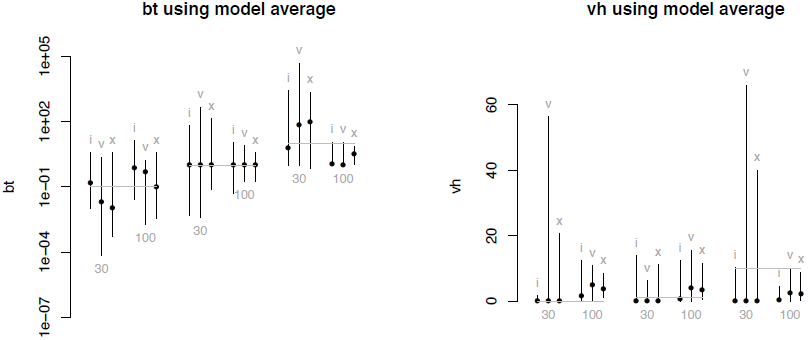
Shown are estimates for *β* (left) and *v*_*H*_ (right) under the simulations. Gray bar shows the true values, dot shows the median model averaged estimate across simulations, and error bars show the range for 95% of the simulations. “i”, “v”, and “x” represent simulations with 1, 5, or 10 hybrid taxa; 30 or 100 refers to the number of non-hybrid taxa. Note *β* estimates often centered on the true value, while *v*_*H*_ was often very wrong.

Even for *β*, however, there is cause for concern. Though the median estimates across simulations were fairly close to the true values, the range across simulation replicates was still quite extreme. For example, for 100 taxon trees with 10 hybrids, the “best case” scenario we examined, if *β* was truly 1.0, indicating no expected bias between the hybrid and the mean of its parents, estimates of beta ranged from 0.13 to 5.14; that is, if we were looking at an organism whose parent species were each 100g, and we used log(mass) as the trait undergoing Brownian motion, the expected mass of the hybrid would be anywhere from 13g (=exp(log(100) + log(0.13)) to 514g according to the estimates. Across the range of the 675 simulations that completed (some with 30 nonhybrid taxa and only one hybrid taxon), where the true value of *β* was 1, the expected mass of a hybrid of two 100g species based on the estimated *β* could be anywhere from 0.00001 g to 10,707,208 g.

### Assessing Model Identifiability through Jointly Estimating Parameters

The shape of the likelihood surface provides the ability to estimate parameter values: if the surface is flat, there is little support for a parameter estimate. In some cases, like the trend parameter for Brownian motion with a trend for coeval taxa, no amount of data is adequate to estimate the parameter: this parameter is formally non-identifiable. There is also a softer definition of identifiability: given a particular dataset, is there enough data to estimate a parameter. We investigated both of these by creating contour maps of the likelihood surface for pairs of parameters under the cichlid and *Nicotiana* data sets. Results are shown in Figure 4. For these empirical datasets, parameters appear distinguishable: there are no ridges in the likelihood surface, even though the confidence intervals are wide. Thus, the parameters are formally identifiable. The large confidence intervals, though, suggest that they can often be practically problematic.

**Figure 4:**
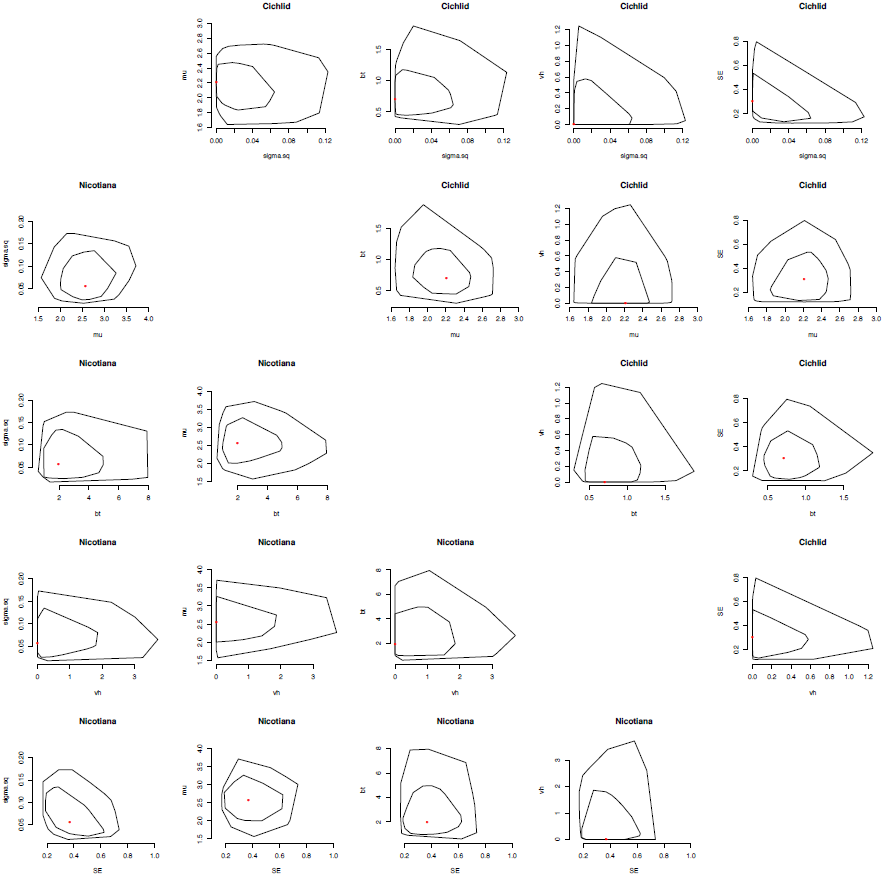
Likelihood surfaces for pairs of traits. Results from cichlids are shown above the diagonal, *Nicotiana* below. The red dot represents the maximum likelihood estimate; the inner circle shows the Δ2 log likelihood unit region, and the outer shows the Δ5 log likelihood unit region. Note the lack of ridges and the wide intervals.

**Figure 5.**
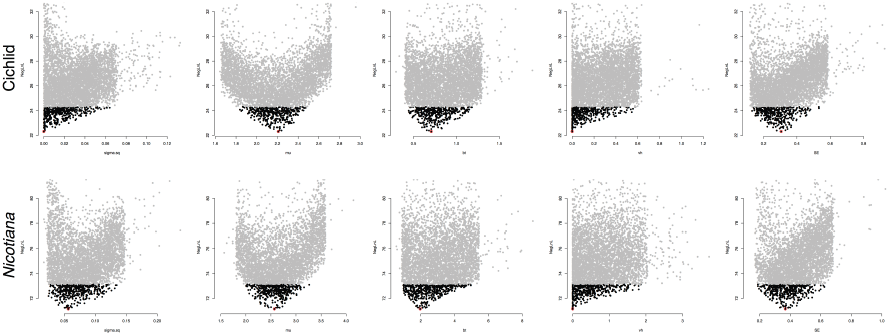
The adaptive confidence intervals for the free parameters of the best model for cichlids (top) and for *Nicotiana* (bottom). In each plot the vertical axis represents the negative log likelihood value and the horizontal axis represents a wide parameter range. The red dot at the lowest y-axis value represents the MLE estimated from the overall search. Sampling works by proposing a set of parameters and estimating the likelihood for this set. This likelihood value is then plotted versus each of the parameters used in that set in a different subplot as a gray or black dots. Dots in black represent the desired likelihood value (taken as those no more than 2 log likelihood units away from the maximum) and the width of the adaptive confidence intervals are measured by the left most and right model black dots.

### Model Selection and Parameter Estimation for the Empirical Data

The best model for cichlids had no burst of variation at hybridization events nor a bias in hybrid size (Table 1). In models where *β* was allowed to vary, the confidence interval included 1, and for *v*_*H*_, the confidence interval (and MLE in all models) included 0, again suggesting lack of evidence for evolutionary speed ups or mean changes with hybridization. Measurement error was estimated to be substantial; in fact, the model estimated no effective Brownian motion at all (*σ*^2^ estimates of 0) with all observed variance being due to just measurement error at the tips. For *Nicotiana*, the best model (but only with 0.39 of the Akaike weight) had *β* as a free parameter and *v*_*H*_ constrained to be 0. There is weak evidence from this model that beta is greater than one (point estimate from best model is 1.97, but CI is 0.89 to 4.52; model averaged estimate is 1.49), suggesting that hybrids have higher success rates as seedlings under drought conditions than do their parents. There is again little evidence for increased variance at hybridization events. Given the tree height and its *σ*^2^ rate, we expect variance at the tips to be 0.50; from measurement error, there is an additional 0.37 variance (both in units of log((seedling survival)^2^)), suggesting meaningful Brownian motion on the tree but still quite important measurement variance.

**TABLE 1:** Model fitting for the cichlid and tobacco data. In this table, NegLogL is the negative log likelihood of the model, *K* is the number of free parameters, ΔAICc is calculated by subtracting the AICc value from the lowest AICc value among the four models, Akaike weight is calculated by *w* = exp(−0.5ΔAICc) and then normalizing the weights. Parameter estimates and their adaptive confidence intervals are reported. The model-averaged parameter estimate 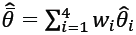 are reported where *w*_*i*_ is the Akaike weight for the *i*th model.

| Dataset | vH parameter | $\beta$ parameter | NegLogL | K | deltaAICc | AkaikeWeight | $\sigma^2$ | $\mu$ | $\beta$ | vH | SE |
| --- | --- | --- | --- | --- | --- | --- | --- | --- | --- | --- | --- |
| Cichlid | fixed | fixed | 23.077 | 3 | 0.000 | 0.524 | 0 (0, 0.095) | 2.152 (1.741, 2.466) | 1 (fixed) | 0 (fixed) | 0.324 (0.091, 0.567) |
|  | free | fixed | 23.077 | 4 | 2.775 | 0.131 | 0 (0, 0.09) | 2.152 (1.759, 2.418) | 1 (fixed) | 0 (0, 0.989) | 0.323 (0.112, 0.556) |
|  | fixed | free | 22.304 | 4 | 1.230 | 0.283 | 0 (0, 0.076) | 2.216 (1.813, 2.501) | 0.708 (0.434, 1.205) | 0 (fixed) | 0.306 (0.112, 0.536) |
|  | free | free | 22.307 | 5 | 4.275 | 0.062 | 0 (0, 0.06) | 2.208 (1.848, 2.482) | 0.707 (0.424, 1.225) | 0 (0, 0.545) | 0.306 (0.132, 0.497) |
| Cichlid model average |  |  |  |  |  |  | 0 (0, 0.087) | 2.174 (1.77, 2.471) | 0.899 (0.804, 1.072) | 0 (0, 0.163) | 0.317 (0.102, 0.552) |
| *Nicotiana* | fixed | fixed | 72.365 | 3 | 0.134 | 0.365 | 0.048 (0.012, 0.112) | 2.591 (1.955, 3.232) | 1 (fixed) | 0 (fixed) | 0.424 (0.244, 0.767) |
|  | free | fixed | 72.275 | 4 | 2.258 | 0.126 | 0.053 (0.019, 0.128) | 2.581 (1.955, 3.171) | 1 (fixed) | 0.229 (0.152, 5.143) | 0.392 (0.195, 0.713) |
|  | fixed | free | 71.146 | 4 | 0.000 | 0.390 | 0.057 (0.014, 0.149) | 2.569 (1.895, 3.252) | 1.968 (0.889, 4.529) | 0 (fixed) | 0.367 (0.178, 0.681) |
|  | free | free | 71.146 | 5 | 2.391 | 0.118 | 0.056 (0.025, 0.148) | 2.568 (1.916, 3.24) | 1.967 (0.827, 4.286) | 0 (0, 1.969) | 0.368 (0.191, 0.623) |
| *Nicotiana* model average |  |  |  |  |  |  | 0.053 (0.015, 0.133) | 2.578 (1.927, 3.233) | 1.492 (0.936, 2.766) | 0.029 (0.019, 0.882) | 0.391 (0.206, 0.71) |

## DISCUSSION

Our new approach allows analysis of trait data on a phylogenetic network. We can now use comparative methods on general networks rather than just trees and it works well for estimating general Brownian motion parameters such as evolutionary rate *σ*^2^ and root state *μ*. There are also hybridization-specific parameters where it performs variably. The method can estimate hybridization bias *β*, the consistent increase or decrease in trait value upon hybridization that may lead to hybrid vigor or outbreeding depression, though with substantial uncertainty. It performs surprisingly poorly for estimating a burst of variance associated with hybridization *v*_*H*_; we would recommend caution interpreting any estimates of *v*_*H*_. Unfortunately, this may be the most biologically interesting question.

Our empirical results for the cichlid dataset do not provide great biological insights, other than 1) the feasibility of running a model given a network and 2) quite extensive measurement uncertainty in the body length measurements. This latter could reflect real measurement uncertainty (fish have indeterminate growth (Dutta 1994)), so the notion of a true species mean for this trait is problematic) but errors in the tree topology or branch lengths would tend to result in this appearing as measurement error in this model, as well. *Nicotiana* also had substantial measurement error, but not enough to wipe out the phylogenetic history. The results suggest that hybrids perform better in droughts than their parent species, though this is not statistically significant given the confidence interval. However, it might point the way to further studies about drought tolerance, an area that will be of increasing importance.

Several approaches have been proposed for inferring different rates along the branch for a given phylogenetic tree (McPeek, 1991; O’Meara et al., 2006; Revell, 2008; and Beaulieu et al. 2012), and it would be useful to extend our work to allow for this heterogeneity. Another possible extension is to use a more parameter-rich Markovian process. The model can be extended to allow trait evolution following the OU process (Hansen 1997; Butler and King 2004; Beaulieu et al. 2012). Putting the approach in a Bayesian context is also possible. In this case, parameters of multiple selective regimes, multiple rates of variation, and multiple rates of constraining forces could be embedded in the model. Developing a more complex model of this type could be very useful when analyzing fairly large data sets of hundreds of species or more, where heterogeneity is expected and there may be power to provide estimates for many parameters.

In nature, approximately 10% of animal species and 25% of plant species hybridize (Mallet 2005, 2007), suggesting that there is widespread gene flow between “species.” Some of this gene flow may lead to hybrid speciation in the manner assumed in our method, and hybrid speciation is widely suspected in many groups (Arnold 1996, Welch and Riesberg 2002). However, even in the absence of hybrids formed from two distinct parent species, such ongoing gene flow suggests a need for a network metaphor, as suggested by Morrison (2014). Our method cannot currently deal with this sort of gene flow: we represent the hybrid as being the result of a single event between two parent lineages (though we do allow for one or both parents to be missing from the tree, making the event appear as if it is going forward in time from the nearest sampled relative(s)). Gene flow over continuous time periods is thus not modeled yet, though it would be a basic extension.

This approach, especially the creation of the modified variance covariance matrix given hybridization and the potential for a modified matrix of expected species values, could form the core for multivariate approaches, in the same way the traditional Brownian motion tree model lies at the heart of methods as various as PGLS (Martins and Hansen 1997), PGLM (Ives and Helmus 2011), independent contrasts (Felsenstein 1985, 2008), phylogenetic linear regression (Ho and Ané 2014) and more. It is also trivial to use the existing approach to estimate ancestral states on a network. While network inference methods are advancing, it is important to make sure that comparative methods using these networks keep pace, of which this work is a start that we hope can be built upon.

## ACKNOWLEDGEMENTS

We thank Luke Harmon and an anonymous reviewer for their suggestions that improves this manuscript. We also thank the National Institute for Biological and Mathematical Synthesis (NIMBioS), an Institute sponsored by the National Science Foundation, the U.S. Department of Homeland Security, and the U.S. Department of Agriculture through NSF Award #EF-0832858, with additional support from The University of Tennessee, Knoxville for funds for a postdoc for D-C J and for summer visitor support, as well as a grant to D-C J from National Science Council grant of Taiwan, ROC, NSC-102-2118-M-035-004.

